# Atomistic simulation study reveals transduction of mechanical work generated by ATP hydrolysis onto myosin II functional loops

**DOI:** 10.64898/2026.01.22.700696

**Authors:** Ikuo Kurisaki, Hideo Higuchi, Shigenori Tanaka, Madoka Suzuki

## Abstract

Myosin II is a paradigm of biological molecular energy transducers that convert the chemical energy of ATP, via hydrolysis, into mechanical work with high efficiency in living cells. The physicochemical mechanisms underlying energy conversion by myosin II have been extensively investigated. However, a remaining challenge concerns the initial stage of the ATP hydrolysis cycle, specifically the conversion of ATP-to-ADP:Pi, because of technical difficulties in directly and seamlessly observing atomic trajectories during this early step and the subsequent processes. Here, we simulate the consequences of the chemical reaction by switching force field parameters between reactant and product systems within classical molecular dynamics simulations. We consider two possible ATP hydrolysis mechanisms in which either singly protonated (Pi²⁻) or doubly protonated (Pi⁻) inorganic phosphate is produced. Our results indicate that, in both Pi-generating processes, the kinetic energy supplied by conversion of ATP-to-ADP:Pi increases only transiently, whereas myosin functional loops store several kcal/mol of potential energy. Meanwhile, the amount of potential energy stored in the Pi²⁻-producing reaction is approximately five-fold larger than that in the Pi⁻-producing reaction. Our analysis indicates that this difference emerges when ATP-derived mechanical work is transmitted into the functional loops via rearrangement of intermolecular hydrogen bonds with the hydrolysis products. Notably, these steric interactions remain stably established even when the kinetic energy input associated with ATP-to-ADP:Pi conversion is actively quenched. We therefore propose that transient storage of ATP-derived mechanical work as atomic-scale conformational strain within myosin molecules constitutes a critical step for efficient conversion of ATP chemical energy into mechanical work under conditions of intracellular thermal noise.

**Significance:** Myosin II converts chemical energy released by ATP hydrolysis into mechanical work with exceptionally high efficiency, even in the presence of substantial thermal fluctuations in living systems. It has long been proposed that release of inorganic phosphate (Pi) from myosin II is a critical step for efficient energy transfer. However, this prevailing view has been challenged by recent state-of-the-art experimental observations, necessitating a reexamination of the chemical steps responsible for ATP-derived energy storage. Here, we employ atomistic molecular dynamics simulations specifically designed to capture key aspects of ATP hydrolysis. We demonstrate that ATP hydrolysis increases the potential energy of myosin II functional loops while Pi remains bound, and that this increase occurs independently of heat release associated with the reaction. Together, these findings identify an alternative mechanism for energy retention and emphasize the intrinsic robustness of myosin II as a highly efficient molecular motor.

## Introduction

Biological nanomachines are physicochemical entities of biochemical processes that sustain living cells (1–6). They hydrolyze ATP or GTP to yield chemical energy of approximately 12 kcal/mol and subsequently perform mechanical work for their functional expression. Because the chemical energy of nucleotides is utilized with remarkably high efficiency, for example exceeding 50% for myosin II and kinesin (7), substantial attention has focused on how their mechanical designs differ fundamentally from those of macroscopic heat engines, which convert thermal heat into mechanical work. Under thermal fluctuations, atomic heat dissipates rapidly into the surrounding environment on picosecond timescales (8–10), far shorter than the timescales required for functional expression of biological nanomachines, which are typically on the order of microseconds or longer. Despite this disparity, biological nanomachines employ mechanisms that retain ATP/GTP-derived chemical energy internally across subsequent chemical steps, thereby preserving this energy until the appropriate timing of functional expression.

Myosin is a paradigmatic nanomachine that has been extensively studied (11–15). This dimeric ATPase generates mechanical force to slide actin filaments through specific steps of the ATP hydrolysis cycle in each ATPase domain. First, *apo* myosin (1. rigor step) binds ATP and detaches from the actin filament (2. prerecovery step), leading to repositioning of the myosin lever arm (3. postrecovery step). The ATP hydrolysis reaction then promotes myosin binding to actin filaments (4. prepowerstroke). Myosin generates mechanical force accompanied by release of the hydrolysis products, inorganic phosphate (Pi), followed by ADP (5. powerstroke step) (16, 17) and subsequently returns to the *apo* myosin state. It has been suggested that the physicochemical origin of myosin’s unusually high energy-conversion efficiency is coupled to Pi release from the ATP hydrolysis reaction site (16, 17). In this scenario, ATP-derived energy is retained inside the reaction site in the form of the reactant molecule Pi until the energy is employed for function, while preventing loss of the generated energy through nondirectional thermal dissipation.

Meanwhile, recent studies have challenged this conventional scenario. According to fluorescence resonance energy transfer (FRET) methods (18, 19), single-molecule laser trap assays (20), and electron microscopy-resolved structures (21), myosin’s force-generating powerstroke can precede Pi release. Consistent with these experimental observations, a theoretical study employing a kinetic model of myosin powerstroke generation reported that slower Pi release results in greater mechanical force generation (22). These findings suggest the existence of alternative chemical steps involved in retention of ATP-derived mechanical energy, beyond Pi binding at the myosin II catalytic site. To examine whether such an alternative mechanism exists for storing ATP-derived energy, and to clarify its nature if present, the most direct approach is to track atomistic molecular dynamics (MD) trajectories that undergo ATP hydrolysis. However, seamless monitoring of ATP hydrolysis from its onset through subsequent chemical steps in myosin at atomic resolution remains challenging, even with state-of-the-art experimental techniques that have been applied to other proteins (23–28).

Here, we address this technical challenge by employing atomistic MD simulations with effective treatments of chemical change, specifically ATP–ADP:Pi conversion, an approach that has been used successfully to provide physicochemical insights into dynamic reaction processes in complex systems (8, 10, 29, 30). This simulation approach, referred to hereafter as switch force field MD (SF2MD), represents the effects of chemical reactions by switching atomic force field parameters between reactant and product systems. Our previous study has demonstrated the practical performance of the SF2MD framework in examining a paradigmatic GTPase, Rat Sarcoma (Ras); in that study we challenged a conventional view of the mechanisms underlying ATP/GTP-generated mechanical work (31) and provided new insight into mechanical work generation via GTP hydrolysis (8).

We employed *Dictyostelium discoideum* myosin II subfragment-1 (**Figure 1A** and **B**) as a model system. Among myosin species that have been suggested to generate force prior to Pi release (18–20, 32–34), the ATP hydrolysis mechanisms of *D. discoideum* myosin II, simply referred to as Myosin hereafter, have been extensively investigated using quantum mechanics/molecular mechanics (QM/MM) simulations (35–41). Since cytosolic ATP predominantly exists as the Mg²⁺-complexed form MgATP²⁻ under physiological conditions (42, 43), prior QM/MM studies (37, 40) have considered two alternative microscopic protonation states of inorganic phosphate generated at the myosin active site, HPO₄²⁻ (Pi²⁻) and H₂PO₄⁻ (Pi⁻). However, it remains unresolved whether one of these species predominates in the prepowerstroke state or whether both species coexist in a defined proportion. Accordingly, we explicitly examined each ATP hydrolysis product in our simulations. Using SF2MD-derived atomistic MD trajectories that focus on the early stage of the ATP hydrolysis cycle, namely the conversion of ATP-to-ADP:Pi, we identify the essential chemical step at which atomic structures store ATP-generated mechanical work prior to Pi release from Myosin. These results enable discussion of the possible role of structure-derived energy storage in Myosin’s chemo-mechanical energy conversion with remarkably high efficiency.

**Figure 1.**
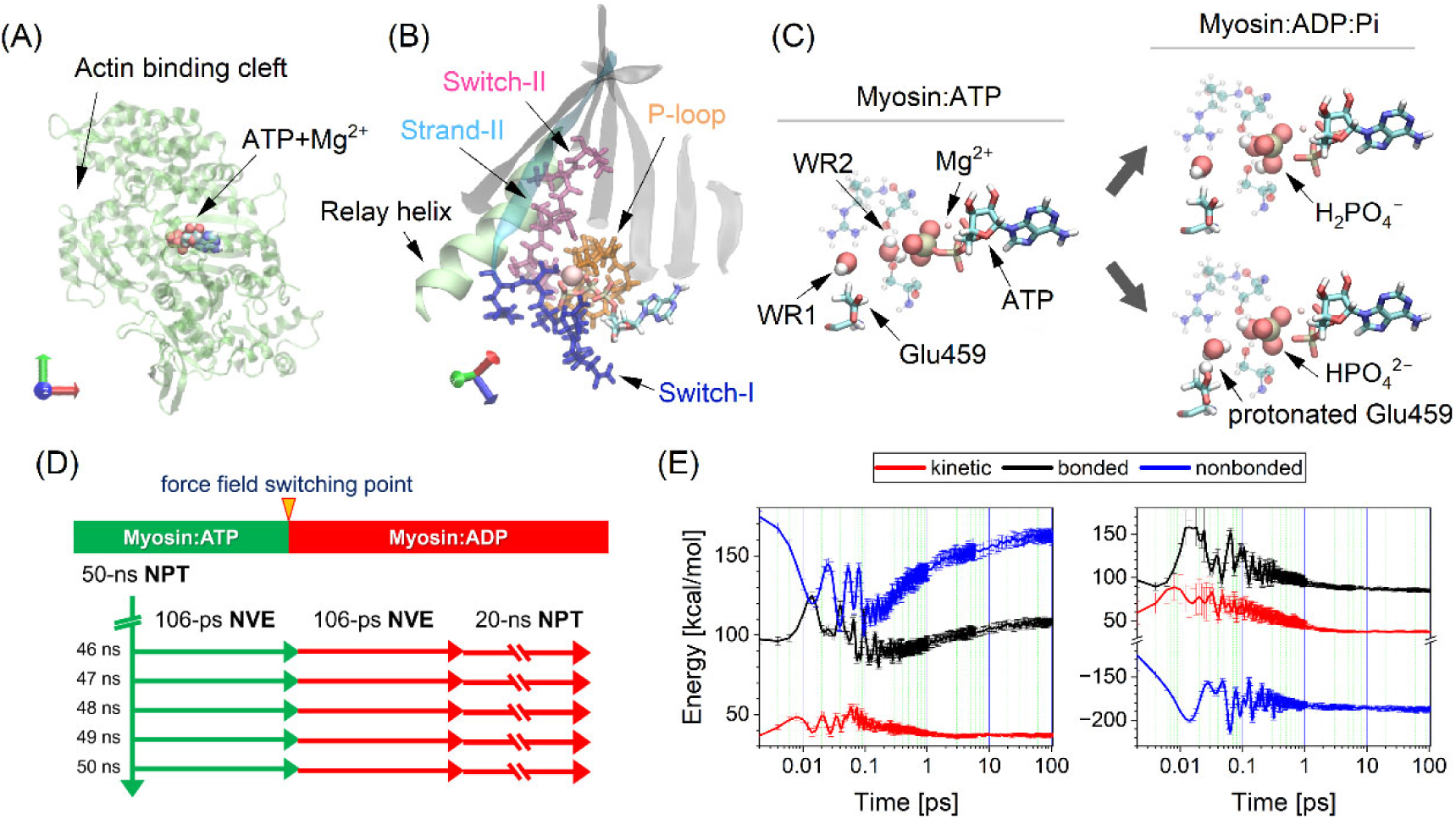
Tertiary structure of subfragment 1 of *D. discoideum* myosin II (hereafter Myosin). PDB entry: 1VOM. (A) Overall structure. Myosin and ATP are illustrated as a green ribbon and a van der Waals sphere, respectively. (B) Enlarged view of the ATP-binding domain. Orange, blue, and magenta sticks denote the phosphate-binding loop (P-loop), Switch-I, and Switch-II, respectively. Strand II and the remaining six strands of the seven-strand sheet are shown in transparent cyan and gray, respectively. ATP and Mg²⁺ are illustrated as a stick and a pink sphere, respectively. A portion of the relay helix (residues 466–481) is shown as a green ribbon. (C) Chemical conversion from the Myosin:ATP system (left) to the Myosin:ADP:H₂PO₄⁻ system (top right) or the Myosin:ADP:HPO₄²⁻ system (bottom right) using QM/MM simulation. ATP and ADP are depicted as solid sticks. H₂PO₄⁻ (Pi⁻), HPO₄²⁻ (Pi²⁻), and two reactive water molecules (i.e., WR1 and WR2) are shown as van der Waals spheres. The Mg²⁺ ion is shown as a pink sphere. Neighboring residues (i.e., Ser181, Arg236, and Ser238) are displayed as transparent ball-and-stick representations. (D) Schematic of the SF2MD simulation procedure. (E) Time course of mechanical energy changes of ADP:Pi within Myosin. The Pi⁻-generation process (left) and the Pi²⁻-generation process (right) are shown. Kinetic energy and bonded and nonbonded contributions to potential energy are indicated by red, black, and blue lines, respectively. Error bars represent 95% confidence intervals.

## Results

We initiated the study with 50 independent and unbiased 50 ns MD simulations of the *D. discoideum* myosin II–ATP complex (Myosin:ATP, hereafter) system (**Figure 1A** and **B**) under constant particle number, pressure, and temperature (NPT)conditions (**Figure 1D**). Temporal changes in root mean square deviation (RMSd) indicated that the X-ray-resolved Myosin structure reached equilibrium in aqueous solution by 40 ns (**Figure S1**). From the final 5 ns of each of the 50 independent 50 ns NPT-MD simulations, five sets of atomic coordinates together with atomic velocities were extracted at 1 ns intervals. This procedure yielded a total of 250 snapshot structures (50 × 5). Each of these 250 snapshot structures was then simulated for 106 ps under constant particle number, volume, and energy (NVE) conditions (five horizontal green arrows in **Figure 1D**). The atomic coordinates and velocities obtained from these NVE-MD simulations were subsequently used to construct the Myosin:ADP systems. Previous QM/MM studies have consistently reported localization of Mg²⁺ in the vicinity of ATP upon hydrolysis, forming a reactive configuration (37, 40) In agreement with these reports, Mg²⁺ was positioned around Pγ and Pβ of ATP in all 250 snapshot structures prior to ATP–ADP:Pi conversion, as illustrated by the example shown in **Figure 1C** (left).

Next, we performed a QM/MM simulation for each of the 250 snapshot structures and converted ATP and reactive water molecule into ADP with Pi⁻ or ADP with Pi^2^⁻ (representative structures are shown in **Figure 1C**, top right and bottom right, respectively) (37, 40). As expected, chemical conversion failed when reactive water molecules were absent from positions required to generate physicochemically reasonable atomic configurations of Pi⁻ or Pi²⁻, and these structures were excluded from subsequent analyses. As a result, we obtained 169 and 128 sets of snapshot structures corresponding to ADP with Pi⁻ and ADP with Pi²⁻, respectively. The lower success rate observed for ADP with Pi²⁻ relative to ADP with Pi⁻ can be attributed to the more restrictive requirements for Pi²⁻ generation. Specifically, formation of ADP with Pi⁻ requires a single reactive water molecule near ATP (**Figure 1C**, top right), whereas formation of ADP with Pi²⁻ requires two water molecules to be simultaneously positioned appropriately around the reactive site (**Figure 1C**, bottom right).

Extended NVE-MD simulations performed using these Myosin–ADP systems (five horizontal red arrows at the center of **Figure 1D**) were consistent with the postulated post-ATP-hydrolysis mechanical process (31). In this proposed process, repulsive interaction energies acting between ADP and Pi are assumed to be converted into kinetic energy. In accordance with this scenario, energy profiles of ADP:Pi derived from SF2MD trajectories exhibited transient increases in kinetic energy as well as in the bonded contribution to potential energy, defined as the sum of bond, angle, and dihedral energy terms, within the 1 ps time domain in both Pi⁻- and Pi²⁻-generation processes (**Figure 1E**). This energy conversion was examined under NVE conditions, thereby avoiding artificial energy exchange with either the thermostat or the barostat. The observed increases arose from relaxation of the nonbonded contribution to potential energy, defined as the sum of Coulomb and van der Waals energy terms, that is, from resolution of steric interactions between ADP and Pi induced by chemical conversion. The nonbonded energy of ADP:Pi⁻ was lower than that of ADP:Pi²⁻, because the former contains an additional proton on Pi that electrically stabilizes the total potential energy. Taken together, these observations support the physicochemical validity of the microscopic processes simulated here using the SF2MD framework.

### Pi^−^-generated ATP hydrolysis raises the potential energies of functional loops

Next, we explored how ATP hydrolysis-derived energy is transferred to Myosin. To track this process, we first focused on the Pi⁻-generating pathway and examined changes in the mechanical energy of Myosin’s functional regions. Among the 18 regions analyzed (**Table S1**), we identified pronounced changes in three functional loops, namely the P-loop, Switch-I, and Switch-II (**Figure 1B**), which together constitute the ATP hydrolysis reaction center. Corresponding analyses for the remaining 15 regions are summarized in Table S2.

After ATP was converted into ADP and Pi⁻ by effectively simulating the hydrolysis process, the potential energies of the P-loop and Switch-II increased during the 106 ps NVE-MD simulations (right panels of **Figures 2A** and **C**). These increases in potential energy were maintained during the subsequent 20 ns NPT-MD simulations (right panels of **Figures 2A** and **C**). By contrast, the potential energy of Switch-I showed a transient increase within the first 10 ps of the 106 ps NVE-MD simulations and then decreased to a level below the initial value by the end of the NVE-MD simulation (**Figure 2B**, right).

**Figure 2.**
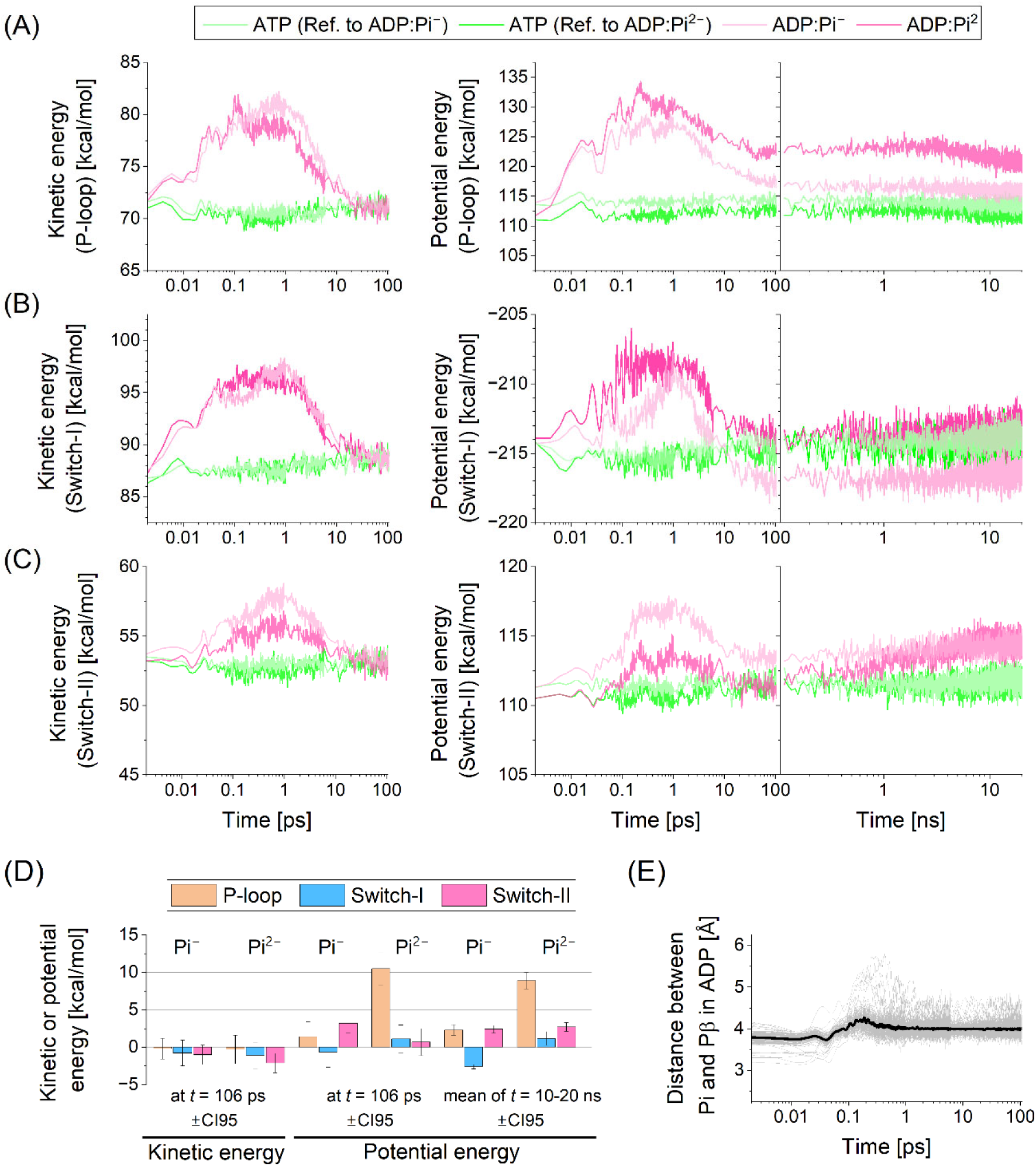
Mechanical energy changes in *D. discoideum* myosin II (Myosin):ADP:H₂PO₄⁻ (Pi⁻) and Myosin:ADP:HPO₄²⁻ (Pi²⁻) systems. (A–C) Time courses of mechanical energy in the P-loop (A), Switch-I (B), and Switch-II (C). Lighter and darker red lines correspond to Myosin:ADP:Pi⁻ and Myosin:ADP:Pi²⁻ systems, respectively, whereas lighter and darker green lines indicate the corresponding Myosin:ATP reference systems. Kinetic energies during the 106 ps NVE-MD simulations are shown in the left panels, while potential energies obtained from the 106 ps NVE-MD simulations followed by the 20 ns NPT-MD simulations are shown in the right panels. Green and pink lines denote the Myosin:ATP and Myosin:ADP:Pi⁻ systems, respectively. (D) Summary of mechanical energies at the indicated time points for the Myosin:ADP:Pi⁻ and Myosin:ADP:Pi²⁻ systems. Error bars represent 95% confidence intervals calculated from 169 and 128 trajectories for the Myosin:ADP:Pi⁻ and Myosin:ADP:Pi²⁻ systems, respectively. (E) Time courses of the distance between the phosphate atom of Pi⁻ and the Pβ atom of ADP during the 106 ps NVE-MD simulations for 169 Myosin:ADP:Pi⁻ trajectories. Black lines indicate average values at each time point across all trajectories, whereas gray lines represent the ensemble of individual MD trajectories.

Meanwhile, the kinetic energies of these functional loops exhibited transient increases during the first several picoseconds. However, consistent with previous nonequilibrium MD simulation studies (8, 10), the kinetic energies rapidly returned to their initial values by the end of the NVE-MD simulations, at 106 ps (left panels of **Figures 2A, B**, and **C**). These observations indicate that ATP-derived energy is converted into potential energy within Myosin. This stored energy persisted during the subsequent 20 ns NPT-MD simulations, implying retention within Myosin until transfer to another functional site accompanied by characteristic conformational changes (**Figure 1D**). This interpretation is further supported by the later analysis of heat-quenched MD simulations.

Such mechanical energy dynamics were not observed in the Myosin:ATP system, as indicated by the green traces in **Figures 2A**, **B**, and **C**. Therefore, we conclude that chemical conversion from ATP-to-ADP:Pi⁻ induces emergent steric interactions between the reaction products and neighboring chemical groups in Myosin, which in turn perform mechanical work on the functional loops and increase their potential energies.

The magnitude of mechanical work required to elevate the potential energies of the functional loops was quantified by analyzing the final 10 ns of the NPT-MD simulations using **Eq. 1** (see Materials and Methods) (**Figure 2D**). For Pi⁻ generation, the potential energies increased by 2.3 ± 0.7 and 2.4 ± 0.5 kcal/mol (mean ± 95% confidence interval) in the P-loop and Switch-II, respectively, whereas the potential energy of Switch-I decreased by 2.6 ± 0.3 kcal/mol. Overall, Pi⁻-generating ATP hydrolysis performed mechanical work of 2.1 ± 1.4 kcal/mol on the three functional loops. Given that ATP/GTP hydrolysis releases free energy of approximately 12 kcal/mol, these results suggest that about 16% of the ATP hydrolysis free energy is converted into mechanical work that increases the potential energies of the functional loops. In addition, this mechanical work generation occurred while the Pi⁻ molecule remained in close proximity to ADP (**Figure 2E**). Thus, mechanical work can be generated prior to Pi release, leading to increased potential energies of Myosin’s functional loops during this initial phase of the ATP hydrolysis cycle.

### The products of Pi^2−^-generated ATP hydrolysis induce a large rise in potential energy

Next, we examined an alternative ATP hydrolysis process in Myosin that accompanies Pi²⁻ generation. During the 106 ps NVE-MD simulations, the kinetic energies of the three functional loops rapidly returned to their equilibrium values within several picoseconds (**Figure 2A**, left). In contrast, the potential energy of the P-loop increased markedly following force field switching and remained elevated throughout the 106 ps simulations (**Figure 2A**, center and right). During the subsequent 20 ns NPT-MD simulations, this potential energy showed a slight decrease but maintained a sustained increase of 8.9 ± 1.1 kcal/mol, as calculated using **Eq. 1** over the final 10 ns of the 20 ns NPT-MD simulations (**Figure 2D**).

Switch-I and Switch-II did not retain the transient increases in potential energy observed during the 106 ps NVE-MD simulations (right panels of **Figure 2B** and **C**, center). However, during the following 20 ns NPT-MD simulations, the potential energies of these regions gradually increased, reaching 1.1 ± 0.9 and 2.7 ± 0.6 kcal/mol in Switch-I and Switch-II, respectively (**Figure 2D**), with values calculated using **Eq. 1** for the final 10 ns of the NPT-MD simulations. In addition to these functional loops, strand II and strand VI within the seven-strand β-sheet also exhibited changes in potential energy of 1.7 ± 0.7 and −1.23 ± 0.24 kcal/mol, respectively (see Table S2, **Figures S2A** and **S3**). Strand II is located near ATP or ADP:Pi but does not form direct atomic contacts with them; however, it is connected to Switch-I along the Myosin amino acid sequence and positioned close to Switch-II (**Figure 1B**). Accordingly, the observed changes in potential energy of strand II are likely triggered indirectly by steric rearrangements of Switch-I and Switch-II induced by conversion of ATP into ADP:Pi²⁻.

Overall, the Pi²⁻-generation process increased the potential energy of Myosin functional loops by a total of 13.1 ± 0.5 kcal/mol (**Figure 2D**). It is therefore plausible that, at this stage of the ATP hydrolysis cycle, a substantial fraction of the free energy released by ATP hydrolysis is captured as increased potential energy within the functional loops and strand II, as suggested by the close correspondence between the observed potential-energy increase and experimentally measured ATP hydrolysis free energy. As observed for Pi⁻ generation, this mechanical work generation also occurs while Pi²⁻ remains bound to ADP (**Figure S4**). Thus, both Pi²⁻- and Pi⁻-generating ATP hydrolysis processes are capable of performing mechanical work on Myosin functional loops and elevating their potential energies prior to Pi release.

### The diversity of inter- and intramolecular hydrogen bond rearrangements is determined by the specific ATP hydrolysis product

Emergent steric interactions induced by ATP–ADP:Pi conversion increase potential energies, although the total magnitude differs between the two ATP hydrolysis processes (**Figure 2D**). To obtain structural insight into these potential energy changes, we analyzed hydrogen bond (H-bond) formation between the functional loops and ATP or ADP:Pi as a hallmark of ATP hydrolysis–induced structural rearrangement. Using the final 10 ns of the NPT-MD trajectories, the numbers of H-bonds and their changes upon ATP hydrolysis were quantified using **Eq. (2)** and **Eq. (1)**, respectively (**Figure 3**). Overall (**Figure 3A** and **B**), both ATP hydrolysis products, ADP with Pi⁻ and ADP with Pi²⁻, form a greater number of intermolecular H-bonds with Myosin than ATP. However, the total number of newly formed H-bonds differs substantially between the two ATP hydrolysis products, as detailed below.

**Figure 3.**
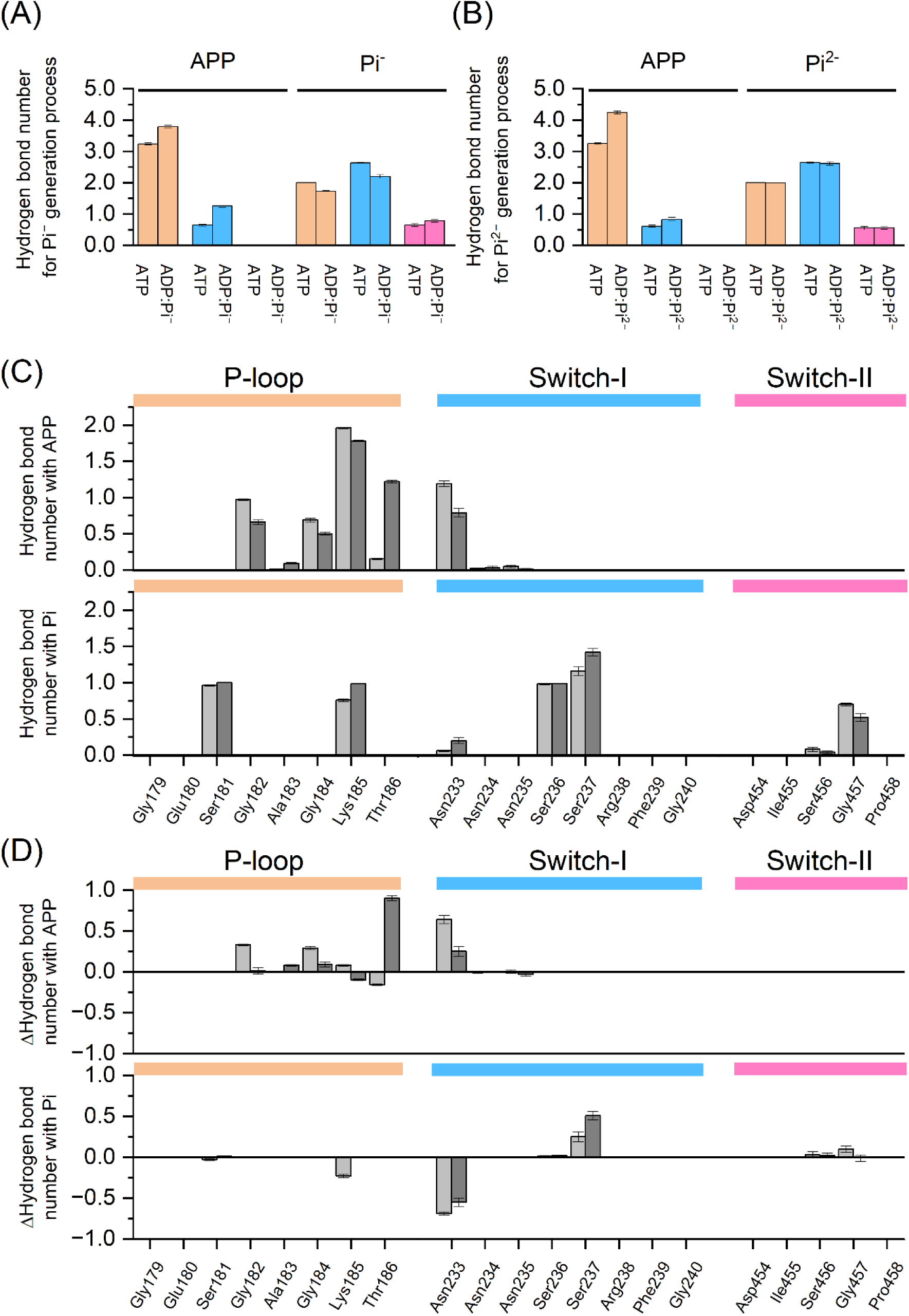
Hydrogen bond (H-bond) rearrangement of functional loops in *D. discoideum* myosin II. Charts are provided for the P-loop, Switch I, and Switch II with ATP or ADP:Pi. Here, Pi⁻ and Pi²⁻ denote H₂PO₄⁻ and HPO₄²⁻, respectively; Pi denotes either Pi⁻ or Pi²⁻. Shown are: (A) H-bond interactions of functional loops with ADP:Pi or ATP during the Pi⁻-generation process. H-bonds associated with the P-loop, Switch I, and Switch II are shown in yellow, cyan, and magenta, respectively. APP denotes ADP in the Myosin:ADP:Pi systems and ATP lacking the PγO₃⁻ group in the Myosin:ATP systems. (B) H-bond interactions of functional loops with ADP:Pi or ATP during the Pi²⁻-generation process. (C) H-bond formation between individual amino acid residues and ADP or Pi. (D) Differences in the number of H-bonds relative to the ATP-bound states, calculated using **Eq. (1)**. In panels (C) and (D), lighter and darker color schemes correspond to the Pi⁻- and Pi²⁻-generation processes, respectively. Error bars indicate 95% confidence intervals. Values shown in panels (A)–(C) were calculated using **Eq. (2)**.

In the Pi⁻-generation process, the number of H-bonds with ADP increases in the P-loop and Switch I by 0.55 and 0.63, respectively (**Figure 3A**). In contrast, H-bonds with Pi⁻ decrease in the P-loop and Switch I and increase in Switch II, with changes of −0.27, −0.43, and 0.13, respectively. As a result, the total intermolecular H-bond number increases across the P-loop, Switch I, and Switch II by 0.28, 0.20, and 0.13, respectively, yielding a summed increase of 0.61 newly formed H-bonds. By comparison, the overall increase in the Pi²⁻-generation process is 1.21. In this case, the number of H-bonds with ADP increases in the P-loop and Switch I by 0.99 and 0.22, respectively (**Figure 3B**), whereas the number of H-bonds with Pi²⁻ remains unchanged in all three loops. In summary, the Pi²⁻-generation process exhibits an approximately two-fold greater increase in intermolecular H-bonds than the Pi⁻-generation process. In general, formation of intermolecular interactions stabilizes the system energetically at the expense of intramolecular interactions. From this qualitative viewpoint, the larger potential energy increase observed in the Pi²⁻-generation process relative to the Pi⁻-generation process can be attributed to differences in the number of newly formed intermolecular H-bonds.

ATP hydrolysis products influence not only the overall increase in H-bonding between Myosin and ADP:Pi but also the underlying microscopic interaction network. The Pi⁻- and Pi²⁻-generation processes share a largely similar set of amino acid residues involved in H-bond formation with ADP. Nevertheless, distinct differences emerge at the level of individual amino acid contributions (**Figure 3C** and **3D**). For instance, the Pi⁻-generation process leads to modest increases in H-bond formation between ADP and Gly182, Gly184, Lys185, and Asn233, whereas the Pi²⁻-generation process produces a pronounced increase in H-bond formation between ADP and Thr186 within the P-loop (**Figure 3D** and **Figure 4A–D**). Examination of strand II in the seven-strand β-sheet, which shows a significant increase in potential energy during the Pi²⁻-generation process (**Figure S2A**), further revealed that Lys241 in strand II approaches Asp453 and Pi²⁻ generated from Pγ of ATP (**Figure S2B**–**D**). Notably, this motion is absent in the Pi⁻-generation process (**Figure S2C** and **D**, lighter red lines).

**Figure 4.**
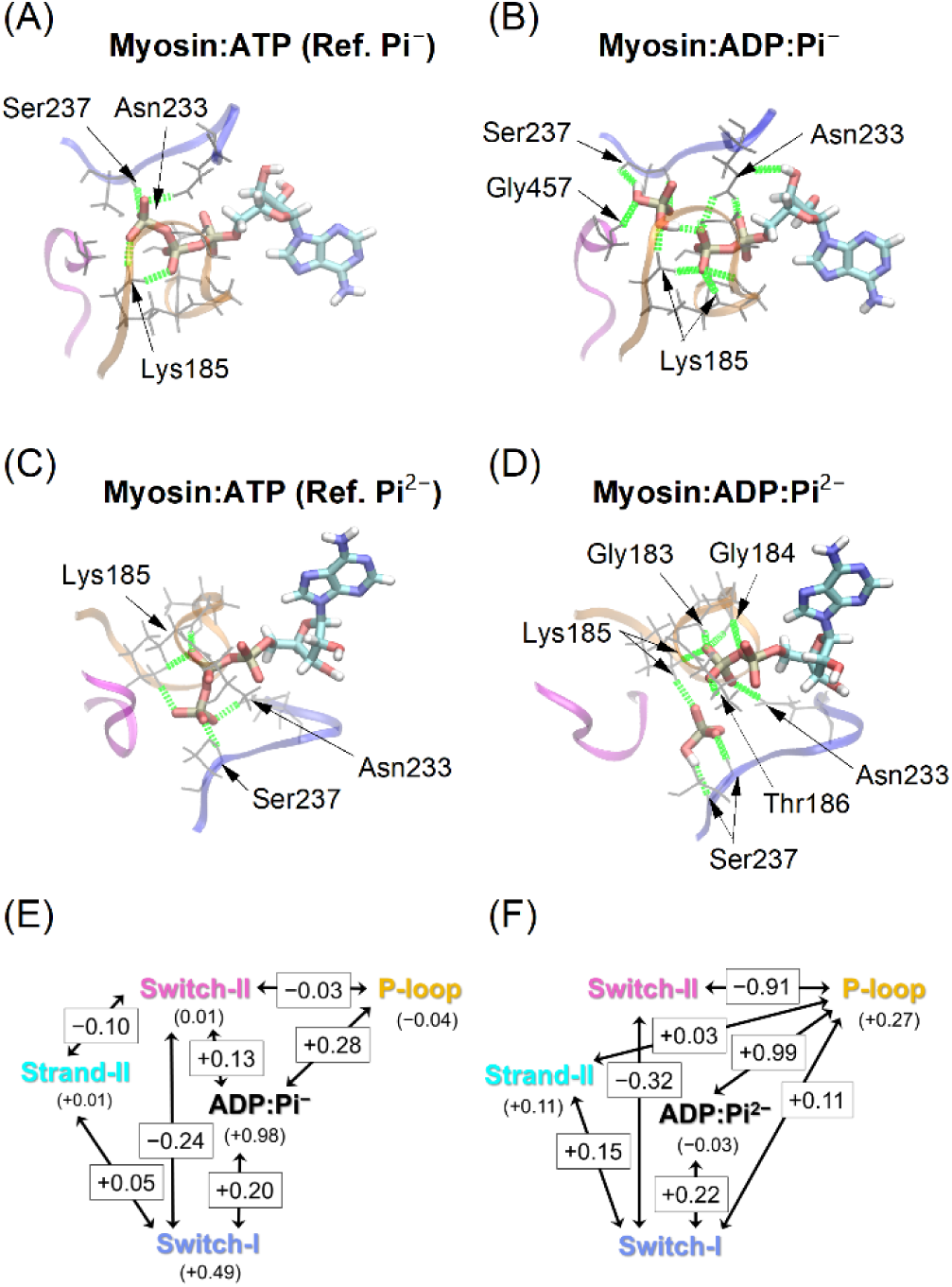
Structural representation of hydrogen bond (H-bond) rearrangements at the ATP-binding site of *D. discoideum* myosin II. Here Pi⁻ and Pi²⁻ denote H₂PO₄⁻ and HPO₄²⁻, respectively. Representative snapshots illustrate changes in H-bond interactions accompanying conversion of ATP-to-ADP:Pi. Panels A and B (C and D) correspond to the ATP-bound and ADP:Pi⁻ (ADP:Pi²⁻) states, respectively. Amino acid residues involved in H-bond rearrangements are highlighted for each ATP hydrolysis product. ATP and ADP:Pi are shown as stick representations. Transparent orange, blue and magenta ribbons indicate the phosphate-binding loop, Switch-I, and Switch-II, respectively. Gray thin sticks represent amino acid residues that form H-bonds with ATP or ADP:Pi. Hydrogen bonds are depicted as green dotted lines. Panels E and F show H-bond networks corresponding to the Pi⁻- and Pi²⁻-generation processes, respectively. Averaged numbers of interdomain/intermolecular and intradomain/intramolecular H-bond formations are indicated in rectangles and parentheses, respectively. Averaged values are omitted where no detectable changes were observed.

Collectively, these observations demonstrate that the post-ATP-hydrolysis state of Myosin is strongly dependent on the identity of the ATP hydrolysis product. Reaction products not only form specific sets of H-bonds with Myosin but also induce rearrangement of the internal H-bond network within the protein (**Figure 4E** and **F**). The patterns of inter- and intramolecular H-bond rearrangement differ between the two Pi-generation processes, indicating that the protonation state of inorganic phosphate plays a central role in determining these differences. Although physicochemical roles of ATP hydrolysis pathways have traditionally been discussed in the context of reaction transition states (44), the present analysis focuses on posthydrolysis structural changes. Accordingly, these results highlight a previously underappreciated role of ATP hydrolysis, namely rearrangement of the hydrogen bond network around functional loops, which enables storage of ATP hydrolysis–generated mechanical work within Myosin.

### Changes in atomistic interactions with Myosin increase the mechanical energy of functional loops

Our final question concerns the origin of the potential energy increases observed in the functional loops. Dissipation of kinetic energy supplied by ATP–ADP:Pi chemical conversion, together with concomitant increases in potential energy, occurs within several picoseconds (**Figure 2**). Recalling Ross’ conjecture (31), one straightforward hypothesis is that the supplied kinetic energy performs mechanical work on the functional loops, thereby increasing their potential energies. However, it remains unclear whether the observed increases in mechanical energy arise through this mechanism. By contrast, our earlier SF2MD study of the Ras–GTP system showed that kinetic energy supply is largely irrelevant to such increases, whereas chemical conversion from GTP to GDP plays an essential role by altering microscopic interactions with Ras. Accordingly, we examined whether kinetic energy constitutes the source of increased potential energy in Myosin by quenching the kinetic energy supplied by ATP–ADP:Pi conversion in SF2MD simulations.

In these simulations, kinetic energy was quenched using a thermostat during the initial 1 ps of the 106 ps NVE-MD simulations (left panels of **Figure S5A**–**C** show results for Pi⁻-generation and left panels of **Figure 5A–C** for Pi²⁻-generation). Under these conditions, transient rises in kinetic energy were effectively suppressed throughout the first 106 ps of the simulations, as evidenced by comparison of the black and red traces in the left panels. In the Pi⁻-generation process, the potential energy of the P-loop increased immediately following force field switching and reached a plateau by the end of the 106 ps simulations (**Figure S5A**, right). Switch-I and Switch-II similarly exhibited decreases and increases in potential energy, respectively (right panels of **Figure S5B** and **C**). These changes persisted during the subsequent 20 ns NPT-MD simulations (right panels of **Figures S5A**–**C**). The final potential energy values were 2.7 ± 0.7, −2.9 ± 0.3, and 2.6 ± 0.5 kcal/mol for the P-loop, Switch-I, and Switch-II, respectively, closely matching those obtained from unbiased NVE-MD simulations of the Myosin:ADP:Pi⁻ system (**Figure 2D**). The Pi²⁻-generation process displayed analogous behavior (**Figure 5**). In this case, the final potential energy values were 9.6 ± 1.0, 1.39 ± 0.8, and 2.7 ± 0.6 kcal/mol for the P-loop, Switch-I, and Switch-II, respectively (**Figure 5D**), again closely corresponding to those derived from the original NPT-MD simulations of the Myosin:ADP:Pi²⁻ system (see **Figure 2D** and related discussion).

**Figure 5.**
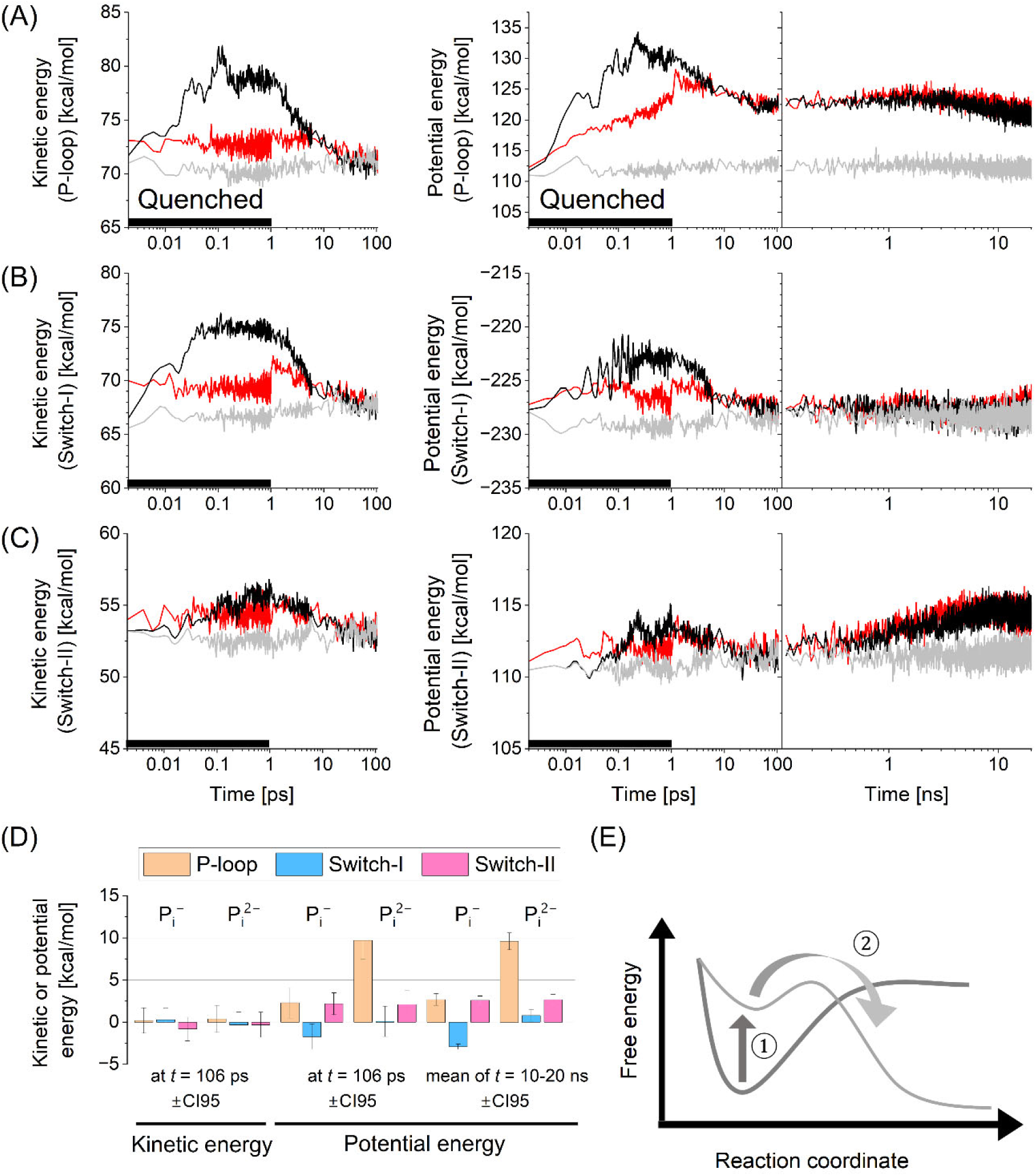
Kinetic energy–quenching simulations in *D. discoideum* myosin II (Myosin):ADP:HPO₄²⁻ (Pi²⁻) system. (A–C) Time courses of mechanical energy in the P-loop (A), Switch-I (B), and Switch-II (C). Kinetic energies obtained from the 106 ps NVE-MD simulations are shown in the left panels, whereas potential energies from the 106 ps NVE-MD simulations followed by the 20 ns NPT-MD simulations are shown in the right panels. Red, black, and gray lines denote quenched, normal, and reference simulations, respectively, for the Myosin:ADP:Pi²⁻ system, where the latter two are taken from Figure 2. Baseline shifts in kinetic energy observed during quenching were caused by use of a thermostat set to 300 K. (D) Summary of mechanical energies at the indicated time points for the Myosin:ADP:H₂PO₄⁻ (Pi⁻) and Myosin:ADP:Pi²⁻ systems. (E) Mechanism of Myosin functional expression suggested by the present study, providing an atomistic basis for integrating Brownian rectifier and deterministic lever-arm models. First, ATP is hydrolyzed to store mechanical energy by increasing potential energy (①). Subsequently, this potential energy is released to perform mechanical work by descending the energy gradient ( ②). Error bars indicate 95% confidence intervals.

In conclusion, ATP hydrolysis performs mechanical work on the functional loops, increasing their potential energies in both Pi⁻- and Pi²⁻-generation processes, even when atomic heat is quenched. In other words, the transient kinetic energy increase driven by ATP–ADP:Pi conversion is largely irrelevant to the mechanical work executed on the loops. These findings indicate that mechanical energy generation in the Myosin system arises from structural relaxation following atomistic interaction changes, or equivalently from the switched potential energy surface, rather than from transient kinetic energy transfer (to be discussed later; cf. **Figure 5E**).

## Discussion

It has long been an open question how biological nanomachines convert the chemical energy of ATP into mechanical work with sufficiently high efficiency to express their function. In the present SF2MD study, we investigated the microscopic mechanism by which ATP-derived energy is efficiently stored in Myosin at the prepowerstroke step, and identified a previously unrecognized mechanism operating in the initial phase of the ATP hydrolysis cycle. Specifically, ATP–ADP:Pi conversion via hydrolysis performs mechanical work on Myosin’s functional loops, where ATP-derived energy is retained while Pi remains bound at the active site. Taken together, these observations suggest that the functional loops constitute the ATP hydrolysis reaction center of myosin and contribute to retaining mechanical energy generated during the early phase of the ATPase cycle. Among these loops, Switch II has been implicated in ATPase activation and in mechanically coupling ATP binding to relay-helix motion (17, 45) (see **Figure 1B** for the positional relationship between Switch II and the relay helix). Taken together, our results indicate that Switch II fulfills multiple functional roles across distinct stages of the myosin ATP hydrolysis cycle. We further found that ADP forms more H-bonds with Myosin than ATP, except at the PγO₃ group, indicating strengthened Myosin–ADP interactions. In addition, rearrangement of H-bond network involving the P-loop and Switch I was observed (**Figure 3A,B**), which may facilitate conformational transitions between pre- and postpowerstroke states. These findings support the view proposed by Moretto *et al* that the powerstroke proceeds from the ADP-bound myosin state (46).

The present results further demonstrate that ATP hydrolysis stably performs mechanical work on the functional loops and that the resulting mechanical energies are retained even when kinetic energy associated with chemical conversion is quenched (**Figures 5** and **S5**). Mechanical work generated by ATP hydrolysis originates from the atomistic change in chemical species from ATP-to-ADP:Pi, which reshapes the potential energy surface of the Myosin system. Consequently, Myosin can robustly express its function independently of transient kinetic energy supply or atomic heat dissipation. In this system, mechanical work generated by emergent steric interactions formed during ATP–ADP:Pi conversion is stored as mechanical energy within the functional loops. It is further suggested that this stored energy may subsequently be transferred to Myosin’s functional sites and contribute to the powerstroke step (**Figure 5E**). This design principle enables proper functional expression of ATPases and GTPases under intracellular thermal fluctuations. We envisage that the present study clarifies the atomistic basis underlying classical physical models of biological nanomachines, including Brownian rectifier and deterministic lever-arm, that is, large downhill free-energy gradient, models (further discussion is provided in the Supplemental Discussion in the Supporting Information). We previously identified a similar mode of mechanical work generation in the GTPase Ras (8), and the present findings in Myosin suggest that the role of ATP/GTP hydrolysis in mechanical work generation is both universal and diverse, as discussed further in the Supplemental Discussion in Supporting Information.

In conclusion, this study elucidates the atomistic mechanisms underlying mechanical work generation during the initial phase of Myosin’s ATP hydrolysis cycle. By analyzing effective ATP hydrolysis simulations within the SF2MD framework, we show that mechanical work derived from ATP hydrolysis manifests as changes in Myosin’s potential energies, and that the high efficiency of chemo-mechanical energy conversion originates microscopically from emergent steric interactions between reaction products and neighboring chemical groups created by ATP hydrolysis. Consideration of this robust mechanism for mechanical work generation provides valuable insight into universal design principles of biological nanomachines and informs efficient design strategies for artificial molecular machines.

## Materials and Methods

### Construction of Myosin:ATP system

We used the X-ray-resolved Myosin structure (PDB entry: 1VOM (47)) to construct atomic coordinates for the Myosin:ATP system. The Myosin:ATP complex was solvated in a rectangular simulation box containing 58947 water molecules, and the system was electrically neutralized by addition of seven K⁺ ions. Interatomic forces were calculated using the AMBER ff14SB force field (48) for amino acid residues, the TIP3P water model (49, 50) for solvent molecules, and JC ion parameters adjusted for the TIP3P water model (51, 52) for ions. Force field parameters for ATP/ADP and Mg²⁺ were taken from those developed by Meagher (53) and Allnér (54), respectively. Empirical force fields for singly protonated (HPO₄²⁻, Pi²⁻) and doubly protonated (H₂PO₄⁻, Pi⁻) inorganic phosphate, which are products of ATP hydrolysis discussed in earlier QM/MM studies (37, 40), were generated using RESP charge calculations at the Hartree–Fock/6-31+G* level of theory with Gaussian09 (55) (see **Figure S6** for details), together with the general AMBER force field 2 (56). Additional methodological details are provided in the Supplementary Methods in the Supporting Information.

All simulations were carried out under periodic boundary conditions. Electrostatic interactions were treated using the Particle Mesh Ewald method, with a real-space cutoff of 9 Å. Vibrational motions involving hydrogen atoms were constrained using the SHAKE algorithm. Each MD simulation was performed with Amber 22 (57) using the GPU-accelerated PMEMD module based on the SPFP algorithm (58) within NVIDIA GeForce RTX 3090 hardware.

### MD simulation procedures

Here, we outline the MD protocols used in this study (illustrated in **Figure 1D**; full details are provided in the Supplementary Methods in the Supporting Information). After temperature and density of the Myosin:ATP system was equilibrated to 300 K and 1 bar, the system was propagated for 50 ns under NPT conditions. Subsequently, sets of atomic coordinates together with atomic velocities were extracted from the 50 ns NPT-MD simulations, and each snapshot structure was further evolved for 106 ps under constant particle number, volume, and temperature conditions. Each snapshot structure at 106 ps was then used to construct the Myosin:ADP:Pi system via the SF2MD protocol, in which reactive atoms were optimized using the above force field parameters and in combination with the PM3 level of theory (59) (see Supplementary Methods in the Supporting Information for details), while the corresponding atomic velocities were carried over. The resulting Myosin:ADP:Pi systems were simulated for 106 ps under NVE conditions, followed by 20 ns NPT-MD simulations. Identical 106 ps NVE-MD simulations and subsequent 20 ns NPT-MD simulations were also performed for the original Myosin:ATP system as a reference, allowing evaluation of the effects of ATP–ADP:Pi conversion on Myosin.

System temperature and pressure were regulated using a Berendsen thermostat (60) with a coupling constant of 5 ps and a Monte Carlo barostat with volume exchange attempted every 100 steps, respectively. Initial atomic velocities were randomly assigned from a Maxwellian distribution corresponding to 0.001 K at the beginning of the NVT-MD simulations. The integration time step was set to 2 fs.

### Kinetic energy quenching simulations

The kinetic energy–quenching simulations were designed on the basis of the 106 ps NVE-MD simulations that followed the force field switching point. During the initial 1 ps interval, the Myosin:ADP:Pi system was coupled to a Langevin thermostat with a collision coefficient of 500 ps⁻¹. The Langevin thermostat regulates the velocity of each atom individually as it fluctuates around a target temperature. In contrast, velocity-scaling methods, such as the Berendsen thermostat, uniformly rescale the velocities of all atoms in the system and are therefore unsuitable for quenching increases in kinetic energy that arise within a limited and specific subset of atoms, as observed in the present case. Detailed workflows of these kinetic energy–quenching MD simulations are provided in the Supporting Information of our earlier study (8).

### Analyses of MD trajectories

Mechanical energy, H-bond formation, interatomic distance, and RMSd were calculated using the cpptraj module as implemented in the AmberTools 22 package (57). For each 50-ns NPT-MD trajectory, we calculated RMSd relative to the X-ray crystallography-derived Myosin structure (47) using the backbone heavy atoms (i.e., Cα, N, C, and O). The geometrical orientation of H-bond formation was as follows: the H-X distance was <3 Å and the X-H-Y angle was >135°, where X, Y, and H denote acceptor, donor, and hydrogen atoms, respectively.

An important effect of chemical conversion was then evaluated by evaluating the difference of *E*(*t*) between the Myosin:ADP system and a reference Myosin:ATP system. Here, *E*(*t*) denotes any physicochemical component, such as potential energy or the number of hydrogen bonds, at a given time point *t*. At this time point, the difference between systems is given by the expression 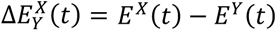. Here, *X* and *Y* are assigned to the Myosin:ADP (M:D) and Myosin:ATP (M:T) systems, respectively. Finally, to characterize the thermal fluctuation of the molecular system, we determined the average of 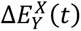 over a specified time interval.

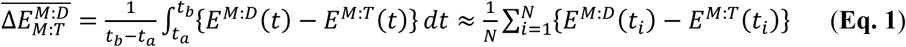

Here, *N* represents the number of snapshots, and *t*_a_ and *t*_b_ correspond to *t*_1_ and *t*_N_ in **Eq. (1)**, respectively.

Similarly, we consider the domain average of *E*(*t*) using the following formula:

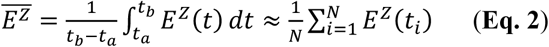

In this equation, *Z* is for either M:D or M:T. The time domain-averaged 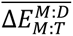 and 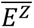 are then further averaged over all simulations for further analyses. Finally, molecular structures were illustrated using visual molecule dynamics (VMD) (61). Error bars show the SE and indicate the 95% confidence interval, unless otherwise stated in the corresponding figure legend.

## Acknowledgments and Funding Sources

This work was supported by JSPS KAKENHI Grant Nos. 22H05053 (to I.K. and M.S.), 22K19273 and 25K02242 (to M.S.). This work was also supported by grants from the Yamada Science Foundation (to M.S.), the Takeda Science Foundation (to M.S.), and the Collaborative Research Program of the Institute for Protein Research, The University of Osaka (Grant Nos. Cra-24-01 and Cra-25-01 to M.S.), and the OU Master Plan Implementation Project of The University of Osaka (to M.S.). Part of the theoretical computations was carried out using the Research Center for Computational Science, Okazaki, Japan (Project: 22-IMS-C014 to I.K. and S.T.), as well as workstations supported by the Core Research for Evolutional Science and Technology Grant Number JPMJCR21F1.

## Competing Interests

The authors declare that they have no competing interests to disclose.

## Supplementary Information

The supplementary information section contains details of atomistic MD simulations, as well as supporting Tables and Figures.

## Data Availability

All data supporting the findings of this study are available from the corresponding author upon reasonable request.

## Notes

### Competing Interest Statement

The authors have declared no competing interest.

