## Supporting Information for "Atomistic simulation study reveals transduction of mechanical work generated by ATP hydrolysis onto myosin II functional loops"

biological nanomachine, energy conversion, molecular motor, molecular dynamics simulation

### Supplementary Methods

#### Technical description of MD simulations

##### System

The X-ray–resolved Myosin structure (PDB entry: 1VOM (1)) contains several unresolved amino acid residues, which were therefore modeled using Swiss-Model (2). The resulting Myosin structure consists of 746 residues, spanning from Asn2 to Arg747. Water molecules present in the crystal structure, as well as the bound $Mg^{2+}$ ion, were retained. Atomic coordinates of the vanadium ion bound to Myosin in the vicinity of ADP were repurposed to define the position of the γ-phosphate in modeling the ATP molecule. The Nε protonation state was assigned to all histidine residues, and all carboxyl groups of aspartate and glutamate residues were set to the deprotonated state. Following these modeling steps, the Myosin:ATP system was solvated in a rectangular box containing 58947 water molecules and electrically neutralized by addition of seven $K^+$ ions.

Empirical force field parameters were prepared for both singly protonated ($HPO_4^{2-}$, $Pi^{2-}$) and doubly protonated ($H_2PO_4^-$, $Pi^-$) inorganic phosphate species. Atomic charges for $Pi^-$ were taken from our previous study on the Ras–GTP–GAP complex (3), whereas those for $Pi^{2-}$ were newly calculated using Gaussian09 (4) (see **Figure S6** for details). The level of theory was selected to ensure consistency with the force field parameters used for ATP and ADP (5). Other force field parameters for these inorganic phosphate species were derived from the general AMBER force field 2 (6). Empirical force fields for the protein, ATP/ADP, water, and ions are publicly available and are described in the main Materials and Methods section, and are therefore not repeated here.

**MD simulations**

We initiated MD simulations using the Myosin:ATP system. Temperature and density of the system were relaxed through the following five-step MD protocol: NVT (0.001 to 1 K, 0.1 ps) → NVT (1 K, 0.1 ps) → NVT (1 to 300 K, 20 ps) → NVT (300 K, 20 ps) → NPT (300 K, 1 bar, 300 ps). During each of these five MD simulations, atomic coordinates other than those of water molecules and $K^+$ ions were restrained by harmonic potentials with a force constant of 100 kcal/mol/Å$^2$ relative to the initial atomic coordinates of each simulation.

The first two NVT-MD simulations were performed with an integration time step of 0.01 fs, whereas the remaining MD simulations employed a time step of 2 fs. The first two NVT-MD simulations were carried out using the Berendsen thermostat (7) with a coupling constant of 0.001 ps, while the subsequent NVT-MD simulations used a Langevin thermostat with a collision coefficient of 1 ps$^{-1}$. In the first and third NVT-MD simulations, the reference temperature was increased linearly over the course of the simulation. The NPT-MD simulation employed a Langevin thermostat with a collision coefficient of 1 ps$^{-1}$ in combination with a Berendsen barostat (7) with a coupling constant of 2 ps. Initial atomic velocities were randomly assigned from a Maxwellian distribution at 0.001 K at the beginning of the first NVT-MD simulation.

Using atomic coordinates obtained from the above relaxation procedure, the Myosin:ATP complex was further structurally equilibrated in aqueous solution through the following seven-step MD protocol: NVT (0.001 to 1 K, 0.1 ps, 10 kcal/mol/Å) → NVT (1 to 300 K, 0.1 ps, 10 kcal/mol/Å) → NVT (300 K, 10 ps, 10 kcal/mol/Å) → NVT

(300 K, 40 ps, 5 kcal/mol/Å) → NVT (300 K, 40 ps, 1 kcal/mol/Å) → NVT (300 K, 40 ps) → NPT (300 K, 1 bar, 50 ns). The final 50 ns NPT-MD simulation was used for RMSd analyses and for subsequent ATP–ADP:Pi conversion.

As above, the first two NVT-MD simulations used a 0.01 fs as integration time step, whereas all remaining simulations used a time step of 2 fs. During the first four steps, Myosin, ATP, and $Mg^{2+}$ were restrained by harmonic potentials with force constants specified in the parentheses above, relative to the initial atomic coordinates. In each NVT-MD simulation, temperature was regulated using a Langevin thermostat with a collision coefficient of 1 $ps^{-1}$. In the final 50 ns NPT-MD simulation, temperature and pressure were controlled using a Berendsen thermostat with a coupling constant of 5 ps and a Monte Carlo barostat with attempts at system volume exchange every 100 steps, respectively. Initial atomic velocities were again randomly assigned from a Maxwellian distribution at 0.001 K. MD trajectories were recorded at 10 ps intervals for subsequent RMSd analyses. The 20 ns NPT-MD simulations performed for the Myosin:ADP:Pi and reference Myosin:ATP systems employed computational settings identical to those used for the 50 ns NPT-MD simulation.

For all MD simulations, translational center-of-mass motion of the entire system was removed every 500 steps to maintain the system near the origin and to prevent overflow of coordinate values in the MD trajectory files. All MD simulations were carried out using the Amber 22 (8) GPU-version PMEMD module based on the SPFP algorism (9) on NVIDIA GeForce RTX 3090 hardware.

**Rearrangement of atomic coordinates for SF2MD**

The hydrolysis reaction accompanies both formation and breakage of chemical bonds among the reactant molecules. Accordingly, we treated this complex rearrangement of chemical bonds with particular care, in contrast to the simpler bond breakage considered in earlier studies (10, 11). To this end, we employed the QM/MM method to simulate chemical conversion from the reactant state (ATP + reactive water molecules) to the product state (ADP + Pi).

As described above, a Myosin:ADP:Pi system was constructed from snapshot structures obtained from each of the preceding 106 ps NVE-MD simulations of the Myosin:ATP system. Positional rearrangements of atoms directly involved in ATP hydrolysis, specifically $P\gamma O_3^-$ in ATP and reactive water molecules, were carried out using QM/MM calculations with the AMBER force field 14 (12) and the PM3 level of theory (13). For the $Pi^{2-}$-generation reaction, Glu459 was additionally allowed to rearrange during the QM/MM simulations, reflecting protonation of its carboxyl group. All QM/MM simulations were performed using the built-in QM/MM interface (14, 15). The QM regions comprised 8 atoms for the $Pi^-$-generation reaction and 13 atoms for the $Pi^{2-}$-generation reaction.

Within this QM/MM procedure, the harmonic potential associated with the interatomic bond between $P\gamma$ and $O_2\beta$ in ATP was removed using ParmEd in Amber22 (8). This treatment is technically required to define the QM region as a closed-shell electronic (8) within simulations performed using the AMBER package.

Based on earlier QM/MM studies of ATP hydrolysis in Myosin (16, 17), water molecule(s) located in the vicinity of Pγ in ATP were selected as reactive water molecules, with a representative configuration shown in **Figure 1C**. We explicitly considered both singly and doubly protonated inorganic phosphate species, $HPO_4^{2-}$ ($Pi^{2-}$) and $H_2PO_4^-$ ($Pi^-$), as products arising from two distinct ATP hydrolysis (16, 17). To generate atomic coordinates for the product states, interatomic distances reported in these earlier QM/MM studies (16, 17) were constrained using harmonic potentials with a force constant of 500 000 kcal/mol/Å$^2$ (**Table S3**). Atom pairs subject to these constraints were selected in accordance with the cited QM/MM studies (16, 17). As noted above, coordinate rearrangements can lead to anomalous increases in kinetic energy if initial atomic velocities are simply propagated. Therefore, to achieve the most seamless practical connection between the preceding and subsequent 106 ps NVE-MD simulations, we applied the minimum necessary set of interatomic constraints to convert ATP and reactive water molecules into ADP together with either $Pi^-$ or $Pi^{2-}$.

Each QM/MM calculation consisted of 100 steps of steepest-descent minimization followed by 400 steps of conjugate-gradient minimization. Atomic coordinates outside the ATP and reactive water molecules region were frozen using the *ibelly* algorithm. Electrostatic interactions were treated using the Particle Mesh Ewald method with a real-space cutoff of 9 Å. Interactions between QM and MM regions were handled using an electrostatic embedding scheme. The total charge of the QM region was set to −2 for both the $Pi^-$-generating and $Pi^{2-}$-generating ATP–ADP:Pi conversion processes.

To clarify the physical validity and scope of the SF2MD approach, we further discuss the underlying separation of timescales and its implications. (18, 19) In the

SF2MD framework, the potential energy surface is switched instantaneously, which may raise concerns regarding proper treatment of the timescale hierarchy among physical processes involved in enzymatic reactions. Following ATP–ADP:Pi conversion, molecular dynamics are described as classical nuclear motion evolving on a newly defined potential energy surface. Electronic rearrangements associated with bond formation and cleavage occur on subfemtosecond timescales ($10^{-16}$–$10^{-15}$ s), at least two orders of magnitude faster than typical nuclear motions ($10^{-14}$–$10^{-13}$ s). With an integration timestep of 2 fs, which is appropriate for Newtonian nuclear dynamics and substantially longer than electronic response times yet shorter than the fastest nuclear vibrations ($\gtrsim$10 fs), the force-field switching in SF2MD can be regarded as an abrupt change in the potential energy surface occurring effectively within a single MD step. From this standpoint, the switching procedure is consistent with the fundamental separation between electronic and nuclear timescales that underpins classical molecular dynamics.

Overall, this treatment is also compatible with an event-based interpretation of enzymatic reactions, as has been emphasized in previous studies of protein dynamics and catalysis. In these studies, slow conformational motions were used to explore the configurational space and occasionally realize reactive configurations, while the chemical conversion itself proceeds as a comparatively rapid event once specific configurations are reached (18, 19). However, SF2MD does not describe how the protein reaches the transition state or resolve reaction kinetics. Instead, it focuses on the nonequilibrium nuclear response that follows product formation by treating the chemical step as an effective, instantaneous perturbation to the system.

The SF2MD approach is therefore most naturally interpreted as a method for probing how chemical conversion induces subsequent structural relaxation, redistribution of energy, and mechanical responses of the surrounding protein environment. In situations where chemical and conformational dynamics are strongly coupled on comparable timescales, or where slow collective motions dominate the reaction process itself, simulation approaches based on continuous reactive potential energy surfaces would be more appropriate. Accordingly, SF2MD should be viewed as a complementary approach within a broader hierarchy of molecular simulation methods, providing a physically grounded and computationally efficient means to analyze postreaction nonequilibrium dynamics under conditions in which a clear separation of timescales can be justified.

### Supplementary Discussion

#### SF2MD simulations provide atomistic insight into biological nanomachines.

The energy conversion mechanisms of biological nanomachines, such as myosin II, have frequently been described using two physical models, the Brownian rectifier and the power stroke (20). The mechanisms embodied in these models are not mutually exclusive, but instead can be combined in an essential manner to achieve high energy conversion efficiency (21). However, the underlying chemical entities associated with these mechanisms are not satisfactorily specified within these theoretical frameworks. Under these circumstances, major advances in single-molecule experiments and structural biology have begun to clarify the mechanistic basis of energy conversion. In parallel, extensive molecular dynamics studies have revealed individual chemical processes, or "steps," that follow ATP hydrolysis (22, 23). The mechanism proposed on the basis of our SF2MD simulations (see **Figure 5E** and related discussion) may provide an atomistic foundation for the flashing Brownian rectifier model, which describes unidirectional motion and associated mechanical work generation through switching of an asymmetric potential energy surface, a well-established physical framework for explaining high energy conversion efficiency in nanomachines (20). In our SF2MD framework, the switching event corresponds to the chemical conversion from the ATP-bound reactant state to the ADP:Pi-bound product state. The microscopic origin of the flashing Brownian rectifier potential energy, at least in part, may therefore arise from changes in the mechanical energies of the functional loops at the moment of ATP–ADP:Pi conversion.

Nevertheless, the present study is limited to 20 ns SF2MD simulations that target the initial phase of Myosin's ATP hydrolysis cycle. By extending SF2MD simulations over timescales sufficient for myosin to undergo global conformational changes, it should become possible to examine the pathways by which energy is transferred from the ATP hydrolysis site to distal functional regions of myosin. Such simulations would enable future investigation of several outstanding questions, including whether mechanical work generated at the hydrolysis site ultimately drives the myosin powerstroke independently of Pi release, whether the magnitude of mechanical work contributing to the powerstroke is comparable to experimentally observed values (24), and how a mechanical interpretation can be developed for the sequential and seamless integration of Brownian rectifier and powerstroke mechanisms (**Figure 5E** ) (20).

#### Generality and diversity of functional loops involved in ATP/GTP hydrolysis energy storage

The role of the P-loop as a site for mechanical energy storage was first reported in a study of mechanical work generation via GTP hydrolysis in Ras GTPase (3). In both Myosin and Ras, the P-loop is positioned in the immediate vicinity of the Pγ atom of ATP and GTP, respectively. This shared positional relationship implies that the P-loop responds sensitively to the generation of Pi in both classes of biological nanomachines. Moreover, the P-loop may therefore serve a conserved function across ATPases and GTPases by storing mechanical work generated through ATP and GTP hydrolysis, respectively.

At the same time, it is informative to consider the diversity of mechanical responses exhibited by molecular nanomachines following ATP or GTP hydrolysis. Here, we focus exclusively on the $Pi^{-}$-generation process, as this pathway is common to both the Myosin and Ras systems. First, Ras utilizes only the P-loop as a site for mechanical work storage (3), whereas Myosin additionally stores mechanical energy in Switch-II (**Figures 2–4**). Second, the magnitude of mechanical work supplied to the P-loop differs between Myosin and Ras, amounting to 2.3 kcal/mol for Myosin (Figure 2D) and approximately 5 kcal/mol for Ras. These differences are likely attributable to variations in the chemical composition of the loops (25), and to their distinct microscopic interactions with other functional domains and with liganded molecules, including ATP and GTP as well as ADP:Pi and GDP:Pi. Taken together, these observations suggest that ATPases and GTPases have diversified their functional roles by modulating how their functional loops respond to ATP and GTP hydrolysis.

#### Supporting Tables

**Table S1**. Functional regions of *Dictyostelium discoideum* myosin II (26, 27).

| Functional region | Residue number range | Functional region † | Residue number range |
|---|---|---|---|
| P-loop | 179–186 | Strand-VII | 116–119 |
| Switch I | 233–239 | Strand-VI | 122–126 |
| Switch II | 454–458 | Strand-IV | 173–178 |
| SH1 helix | 681–690 | Strand-II | 241–247 |
| SH2 helix | 671–680 | Strand-I | 253–261 |
| Strut | 590–593 | Strand-III | 447–453 |
| CM loop | 389–405 | Strand-V | 649–656 |
| Converter domain | 692–747 | | |
| Relay helix | 466–498 | | |
| Relay loop | 499–509 | | |
| Wedge loop | 572–574 | | |

† β-strands that consist of seven strand sheets, listed in the right-side column.

**Table S2**. Summary of changes in potential energies in functional regions, calculated using **Eq. 2**[*]. Values are shown for the last 10-ns time domain of 20-ns NPT-MD simulations [Unit: kcal/mol].

| | $Pi^{-}$-generation | | $Pi^{2-}$-generation | |
|---|---|---|---|---|
| Functional region | Original | Heat quench | Original | Heat quench |
| P-loop | 2.3 ± 0.7 | 2.7 ± 0.7 | 8.9 ± 1.1 | 9.6 ± 1.0 |
| Switch I | 2.6 ± 0.3 | −2.9 ± 0.3 | 1.1 ± 0.9 | 1.39 ± 0.8 |
| Switch II | 2.4 ± 0.5 | 2.6 ± 0.5 | 2.7 ± 0.6 | 2.7 ± 0.6 |
| SH1 helix | 0.5±1.6 | 0.3±1.5 | 1.4±1.4 | –0.7±2.2 |
| SH2 helix[†] | –0.7±1.5 | –0.7±1.4 | 1.8±1.4 | 1.7±1.8 |
| strut | 0.0±0.2 | –0.1±0.2 | –0.3±0.4 | –0.5±0.4 |
| CM loop | 0.6±3.4 | –0.8±3.2 | 2.5±3.6 | 0.2±3.4 |
| Converter[†] | 0.8±8.7 | 2.5±8.9 | 10.4±9.5 | 5.2±10.5 |
| Relay helix[†] | 5.8±3.9 | –0.6±3.7 | 0.1±3.6 | 3.4±4.2 |
| Relay loop | 0.4±1.2 | 1.0±1.3 | 1.0±1.3 | 1.1±1.3 |
| Wedge loop[†] | –0.1±0.2 | 0.0±0.2 | 0.4±0.3 | –0.0±0.3 |
| Strand-I | –0.5±0.5 | –0.3±0.5 | 0.0±0.6 | –0.3±0.5 |
| Strand-II | 0.3±0.8 | –0.2±0.7 | 1.7±0.7 | 1.7±0.9 |
| Strand-III | 0.1±0.1 | 0.0±0.1 | –0.1±0.1 | 0.0±0.1 |
| Strand-IV | 0.2±0.3 | –0.1±0.2 | 0.0±0.3 | –0.3±0.3 |
| Strand-V | 0.2±0.3 | 0.1±0.3 | 0.3±0.3 | 0.0±0.4 |
| Strand-VI | –0.1±0.2 | –0.1±0.2 | −1.2±0.2 | −1.2±0.2 |
| Strand-VII | 0.1±0.2 | –0.1±0.2 | –0.1±0.2 | 0.1±0.2 |

[*]Errors indicate 95% confidence intervals.

[†] Neither the original nor the heat-quenched simulation exhibited a significant difference in potential energies and is therefore not discussed further.

**Table S3**. Pairs of atoms subjected to harmonic potentials in the QM/MM simulations, together with the corresponding equilibrium distances.

| Atom 1 | Atom 2 | Equilibrium distance [Å] |
|---|---|---|
| | $H_2PO_4^-$ generation process | |
| Pγ in ATP | O₃β in ATP | 2.98 |
| Pγ in ATP | O in WR2† | 1.58 |
| O in WR | H in WR2 | 0.98 |
| | $HPO_4^{2-}$ generation process | |
| Pγ in ATP | O₃β in ATP | 2.98 |
| Pγ in ATP | O in WR1† | 1.57 |
| O in WR2 | H in WR1 | 0.98 |
| Oε in Glu459 | H in WR2 | 0.98 |

†WR denotes a reactant water molecule whose oxygen atom is found near the Pγ in ATP and is positioned opposite to the Pγ-$O_3$β bond (see also **Fig. 1C** for WR1 and WR2).

### Supporting Figures

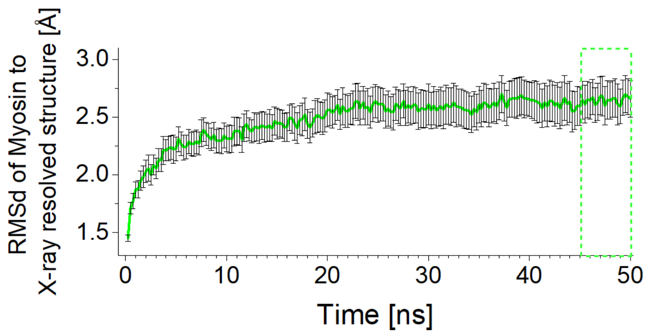


**Figure S1**. **Time course analyses of root mean square deviation (RMSd) for the *D. discoideum* myosin II:ATP system.** The time interval assumed to correspond to RMSd convergence is indicated by a green dotted rectangle. Error bars indicate the 95% confidence intervals. RMSd values were calculated from 50 independent 50-ns NPT-MD trajectories obtained after relaxation of system temperature and density. (300K and 1 bar; see **MD simulation procedure in Supplemental Methods** for details).

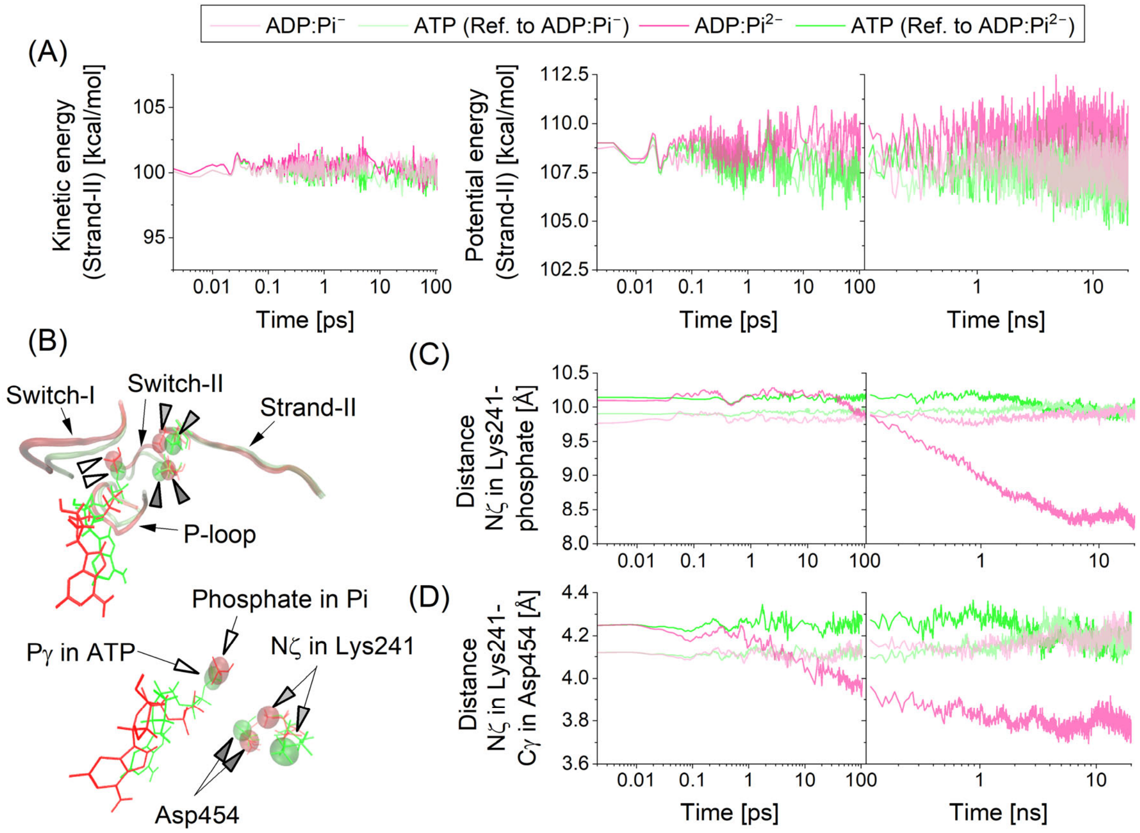


**Figure S2. Effect of ATP–ADP:Pi conversion on Strand II of *D. discoideum* myosin II (hereafter referred to as Myosin).** (A) Time courses of mechanical energy. Kinetic energies during the 106 ps NVE-MD simulation (left) and potential energies during the 106 ps NVE-MD simulation followed by the 20 ns NPT-MD simulation (right). (B) Comparison of atomic positions between the Myosin:ATP and Myosin:ADP:$HPO_4^{2-}$ ($Pi^{2-}$) systems, shown in green and red, respectively. (C) and (D) Time courses of the distance between the phosphate atom of Pi and Nζ of Lys241 and that between Nζ of Lys241 and Cγ of Lys454, respectively. In panels A, C, and D, lighter (darker) red and green lines correspond to the Myosin:ADP:$H_2PO_4^-$ ($Pi^-$) (Myosin:ADP:$HPO_4^{2-}$, $Pi^{2-}$) and the corresponding Myosin:ATP systems, respectively. In panel B, $P_i$ denotes either $P^-$ or $P^{2-}$.

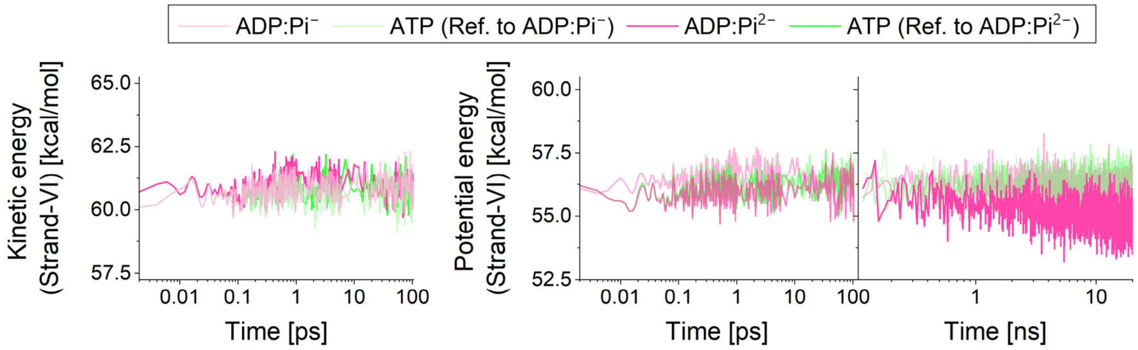


**Figure S3. Effect of ATP–ADP:Pi conversion on Strand VI. Time courses of mechanical energy.** Kinetic energies during the 106 ps NVE-MD simulation (left) and potential energies during the 106 ps NVE-MD simulation followed by the 20 ns NPT-MD simulation (right) are presented.

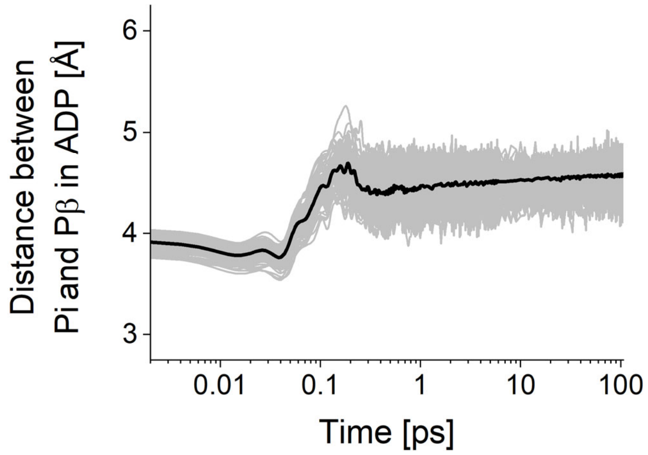


**Figure S4. Time courses of the distance between the phosphate atom of $HPO_4^{2-}$ (Pi) and Pβ of ADP during the 106 ps NVE-MD simulations for 128 Myosin:ADP:Pi trajectories**. Black lines indicate the average distance at each time point across all trajectories, whereas gray lines represent the ensemble of individual MD trajectories.

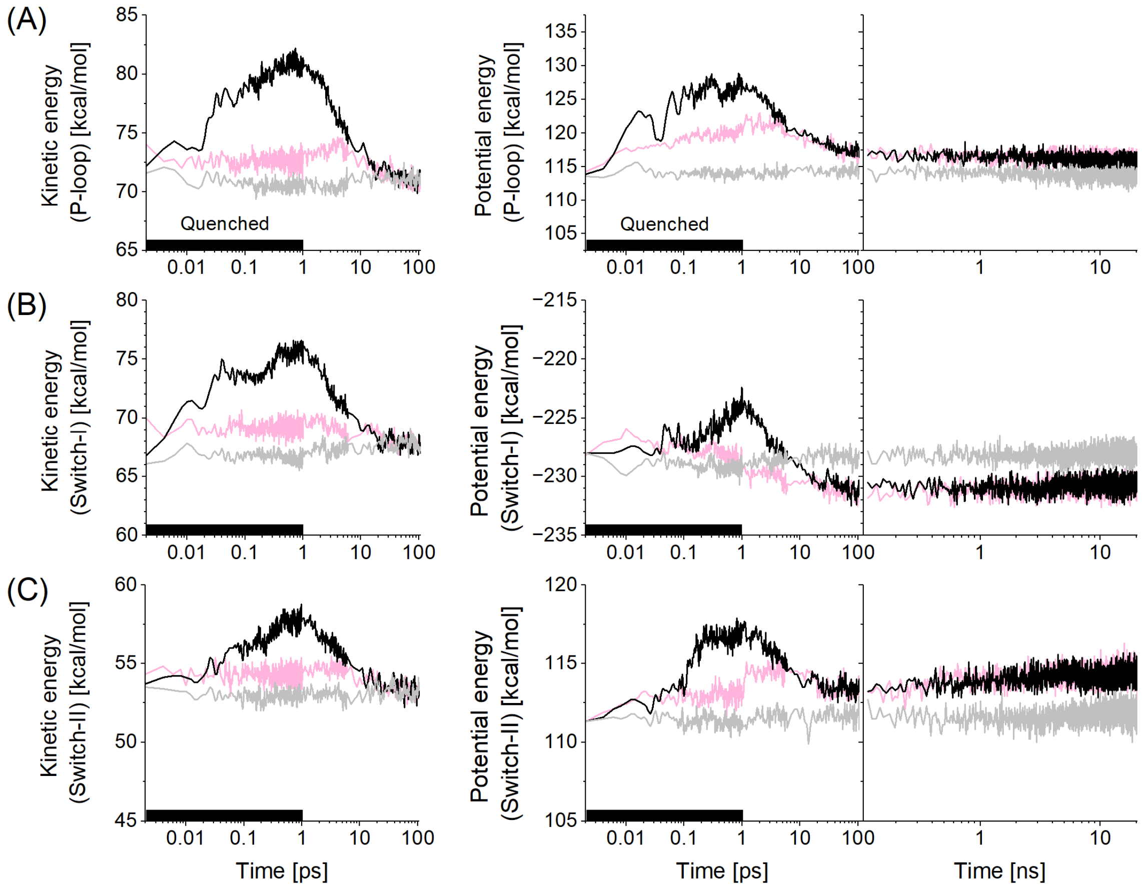


**Figure S5**. **Kinetic energy quenching simulations of the Myosin:ADP:$H_2PO_4^-$ ($Pi^-$) system.** (A) P-loop. (B) Switch-I. (C) Switch-II. Kinetic energies during the 106 ps NVE-MD simulation (left), potential energies during the 106 ps NVE-MD simulation (center), and potential energies during the subsequent 20 ns NPT-MD simulation (right) are shown. Pink, black, and gray lines denote quenched, normal, and reference simulations, respectively, where the latter two are taken from **Figure 2** and correspond to the Myosin:ADP:$Pi^{2-}$ systems.

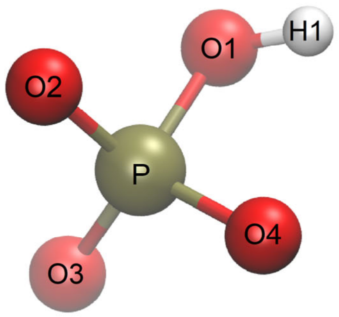


| Atom name† | Atom type† | RESP charge [a.u.] |
|---|---|---|
| O1 | oh | −0.892124 |
| H1 | ho | 0.349940 |
| P1 | p5 | 1.925110 |
| O3 | o | −1.127642 |
| O4 | o | −1.127642 |
| O2 | o | −1.127642 |
| | Total chage | −2 |

**Figure S6**. **Atomic structure of $HPO_4^-$ calculated in vacuum at the Hartree–Fock/6-31G* level of theory.** White, red, and gold spheres represent hydrogen, oxygen, and phosphorus atoms, respectively. Atomic charges for each atom are listed on the right side. Characters shown within the spheres indicate atom names assigned according to the General AMBER Force Field nomenclature.
